# Targeting ductal-endothelial crosstalk alleviate pancreatitis

**DOI:** 10.1101/2024.01.15.575182

**Authors:** Rong-rong Gao, Lan-yue Ma, Jian-wei Chen, Yu-xiang Wang, Yu-yan Li, Zi-yuan Zhou, Zhao-hua Deng, Jing Zhong, Ya-hai Shu, Yang Liu, Qi Chen

## Abstract

Pancreatitis are common gastrointestinal disorders that cause hospitalization with significant morbidity and mortality. The mechanistic pathophysiology of pancreatitis is complicated, which greatly limits the discovery of pharmacological intervention methods. Here, we show that administration of antagonist of Integrin-α5, significantly mitigates the pathological condition of acute pancreatitis. In caerulein-induced acute pancreatitis model, the newly emergent CK19 positive cells are highly vascularized with significant increase of vascular density and endothelial cell number. Single cell RNA sequencing analysis shows ductal and endothelial cells are intimate interacting partners. Pancreatitis dramatically reduce the crosstalk in ductal-endothelial interface but promote the integrin-α5 signaling. Blocking this signaling significantly reduce acinar-to-ductal metaplasia, pathological angiogenesis and restore other abnormal defects induced by caerulein. Our work reveals a therapeutic potential of targeting integrin-α5 as uncharacterized pharmacological method to alleviate the symptom of pancreatitis.

## Introduction

The pancreas is an essential organ for digestion and metabolism, ranking as the second-largest digestive gland(1). Its exocrine portion is comprised of acinar and ductal cells. Acinar cells secrete pancreatic juice, which includes digestive enzymes like amylase, trypsin, and collagenase, to aid in nutrient breakdown(2). Ductal cells, organized in a branched tubular network, transport these enzymes to the duodenum (3).

Pancreatitis, namely pancreatic inflammation, is among one of the most prevalent gastrointestinal disorders that require hospitalization(4). Statistical analysis estimates that the incidence of pancreatitis ranged from 7 to 9 million cases annually worldwide between 2005 and 2015(5). In the United States, 299,150 hospital admissions are diagnosed with pancreatitis, incurring healthcare costs of approximate 8 billion US dollars in 2018(6). The overall mortality rate for acute pancreatitis is reported to be around 20%, which rises sharply to 40% for severe cases(7–9). Severe acute pancreatitis can lead to complications such as sepsis or multiorgan failure(10). But current treatments are mainly supportive care, including intravenous fluid administration, antibiotics to prevent further infection, and pain-killer for patient’s pain management(11). Surgery is utilized to remove damaged pancreatic tissue or obstructive gallstone(12, 13). However, there is an urgent need for pharmacological approach to alleviate the symptoms of pancreatitis.

Although pancreas is a highly vascularized organ, the significance of blood vasculature in pancreatitis is not well-understood. The development of pancreatic vascular network, including branching morphogenesis and arteriovenous specification, is initiated from mid-gestation stage in embryonic mouse(14). During embryonic development of pancreas, hypervascularization is reported to restrain pancreas branching, differentiation and growth(15, 16), suggesting existence of potential interactions between vascular endothelial cells (VEC) and pancreatic resident cells. In other organs, VEC carry out tissue microenvironmental function to crosstalk with corresponding tissue resident cells. For example, in the bone marrow, intestine and brain, VEC form mutualistic relationships with their neighboring hematopoietic stem cell, intestinal stem cell and neural stem cell, respectively(17–20). In several organs, there are close associations between blood vessels and peripheral nerve fibers establishing a crosstalk interface to promote organ regeneration(21–23). However, the interacting partner of blood vessel in the pancreas remain largely unknown.

In this study, we uncover that the branched ductal tubes align parallel with pancreatic blood vessel facilitating the crosstalk between VEC and ductal cells. Targeting pancreatitis-induced Spp-1/Integrin-α5 signaling from ductal cells to VEC by ATN-161 serve as a promising pharmacological approach for mitigating the pathological condition of acute pancreatitis.

## Results

### The CK19^+^ ductal tube parallelly aligned with pancreatic blood vessels

In the emergence of acute pancreatitis, there is a pivotal cellular event in which the pancreatic acinar cells differentiate to CK19^+^ ductal-like cells, a cellular process known as acinar-to-ductal metaplasia (ADM)(2, 24, 25). Given that ADM is a hallmark feature of acute pancreatitis, we examined the spatial distribution of CK19^+^ cells to identify potential cell types that might interact with them.

Co-staining of pan-endothelial cell marker CD31 and CK19 revealed an intimate alignment between pancreatic ductal tubes and the blood vessel network (Figure 1A). This spatial alignment was further confirmed through visualization of pancreatic blood vessels using additional VEC markers including Endomucin (Emcn), vascular endothelial growth factor receptor-2 (VEGFR2), and a genetically modified *Cdh5-mtdTomato-nGFP* (*Cdh5-mTnG*) endothelial reporter mice(17), in which the membrane-anchored tdTomato marked the surface of VEC (Figure 1, B to D and Supplemental Figure 1, A and B).

**Figure 1.**
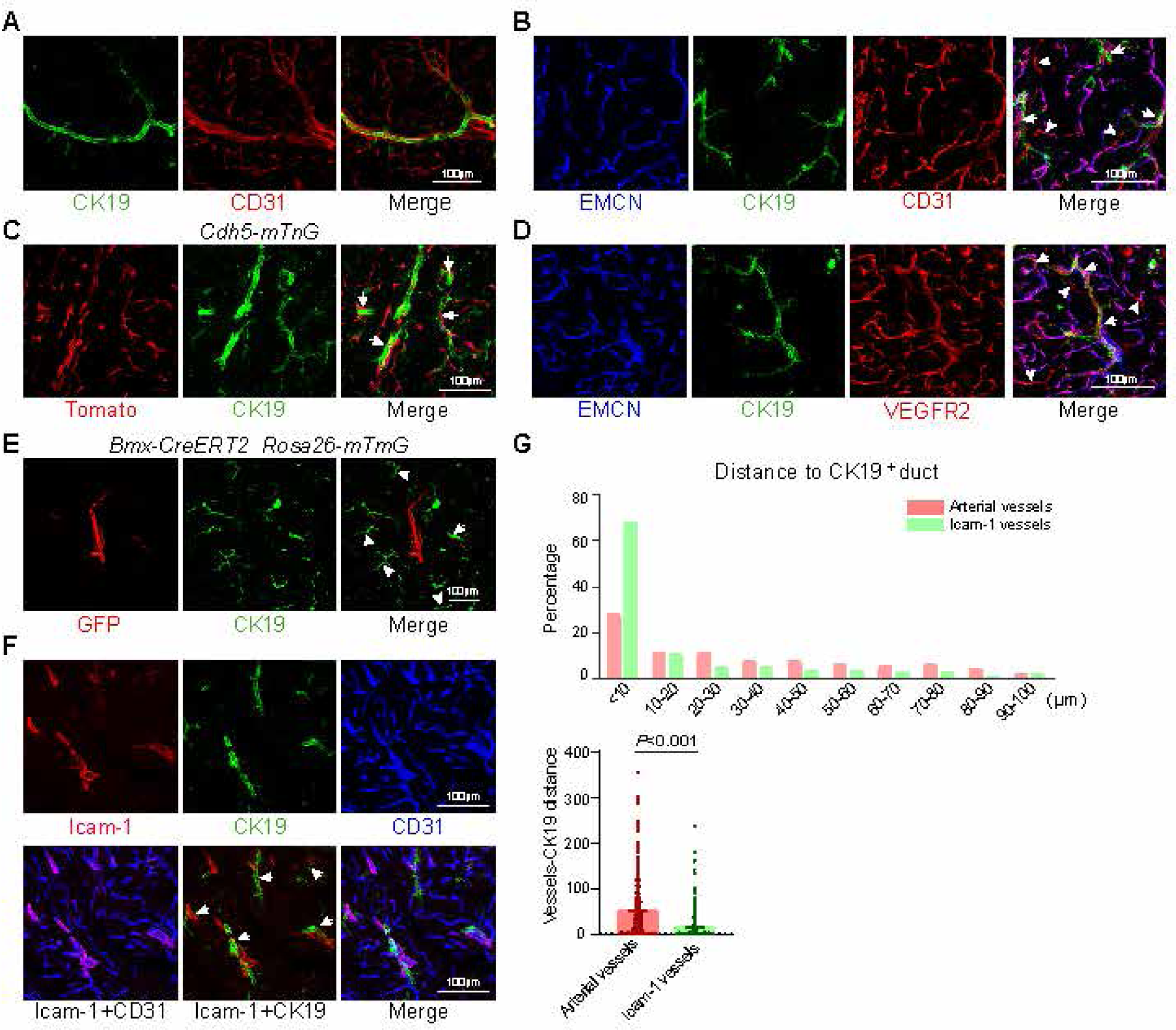
Pancreatic ductal cells were parallelly aligned with blood vessels. (**A**) Representative projected confocal images showing alignment of CK19^+^ ductal cells (green) and CD31^+^ (red) vascular endothelial cells. (**B**) Representative projected confocal images showing co-staining of CK19^+^ ductal cells (green) together with Emcn (blue) and CD31 (red). Arrowheads indicated CD31^+^ Emcn^-^ artery; arrows indicated CK19^+^ ductal-tube associated vessels. (**C**) Representative projected confocal images showing co-staining of CK19^+^ ductal cells (green) together with membrane-tdTomato (red) in *Cdh5-mTnG* endothelial transgenic reporter mice, in which membrane-anchored tdTomato signal labels the surface of VEC. Arrows indicated CK19^+^ ductal-tube associated vessels. (**D**) Representative projected confocal images showing co-staining of CK19^+^ ductal cells (green) together with Emcn (blue) and VEGFR2 (red). Arrowheads indicated VEGFR2^+^ Emcn^-^ artery; arrows indicated CK19^+^ ductal-tube associated vessels. (**E**) Representative projected confocal images showing co-staining of CK19^+^ ductal cells (green) together with membrane GFP signal in *Bmx-CreERT2 Rosa26-mTmG* mice, in which arterial specific *Bmx-CreER* leads to an irreversible switch from constitutive membrane tdTomato expression to membrane GFP expression. Arrow indicated artery closer to ductal cells; arrowheads indicated ductal tube which is not associated with artery. (**F**) Representative projected confocal images showing co-staining of CK19^+^ ductal cells (green) together with CD31 (blue) and Icam-1 (red). Arrows indicated Icam-1^+^ vessels aligned with CK19^+^ ductal cells. (**G**) Accumulative distribution and dot plots to quantitatively compare the distance between arterial and Icam-1^+^ vessels to CK19^+^ ductal cells. Arterial vessels=358; Icam-1^+^ vessels=364. Error bars, mean ± s.e.m. *P* values, Kolmogorov-Smirnov test.

Despite the close spatial relationship between the ductal tube and blood vessels, we observed that CK19^+^ ductal cells aligned with only a subset of pancreatic blood vessels (Figure 1, A to D). We first investigated whether this subset of pancreatic vessels was artery. Similar to other organs, arterial VEC can be identified as CD31^+^Emcn^-^ vessels, tdTomato^+^Emcn^-^ vessels in *Cdh5-mTnG* reporter mice, or more specifically, as membrane-GFP^+^ vessels in *Bmx-CreERT2 Rosa26-mTmG* mice (Figure 1, B to E and arrowheads in Supplemental Figure 1, A to C). Although some arterial VEC were in close proximity to CK19^+^ pancreatic ducts (Arrow, Figure 1E), a large portion of CK19^+^ ductal cells did not align with artery (Arrowheads, Figure 1E). Beyond arteries, we found that Icam-1 could be utilized to distinguish another subpopulation of pancreatic vessels, which were largely separated from arterial VEC (Supplemental Figure 1C). Notably, a substantial proportion of CK19^+^ ductal cells showed proximate location with Icam-1^+^ pancreatic vessel (Figure 1F). Quantitative analysis indicated approximately 70% of Icam-1^+^ vessels were within 10 μm of CK19^+^ ductal cells (Figure 1G). Co-staining of CK19 and Icam-1 at postnatal day (P) 21 showed that this intimate association was already established in the juvenile stage of mouse development (Supplemental Figure 1D). But at P8, we observed that the tubulogenesis of pancreatic duct had not yet finalized, while the blood vessels were differentiating to express Icam-1 (Arrows, Supplemental Figure 1D). This suggests that the parallel alignment between Icam-1^+^ blood vessels and CK19^+^ ductal tube was gradually established during postnatal development of the pancreas.

These findings indicated that pancreatic CK19^+^ cells aligned parallel to blood vessels under physiological condition, suggesting the possibility for these two populations of cells to communicate with each other.

### The caerulein-induced acute pancreatitis was associated with pathological angiogenesis

Next, we investigated whether acute pancreatitis resulted in changes of pancreatic vascular network, because acinar cells can differentiate to CK19^+^ ductal-like cells through ADM during acute pancreatitis. To address this question, we consecutively administrated caerulein (CAE), a molecular ortholog of intestinal hormone cholecystokinin and the most well-established pancreatic inflammation model, to induce acute pancreatitis (Figure 2A) (26, 27). This treatment led to noticeable abnormal tissue morphology, as revealed by Sirius red staining (Figure 2B). An increase in collagen fibers was observed, and the morphology of pancreatic tissue shifted from polygonal to orbicular shapes (Figure 2B).

**Figure 2.**
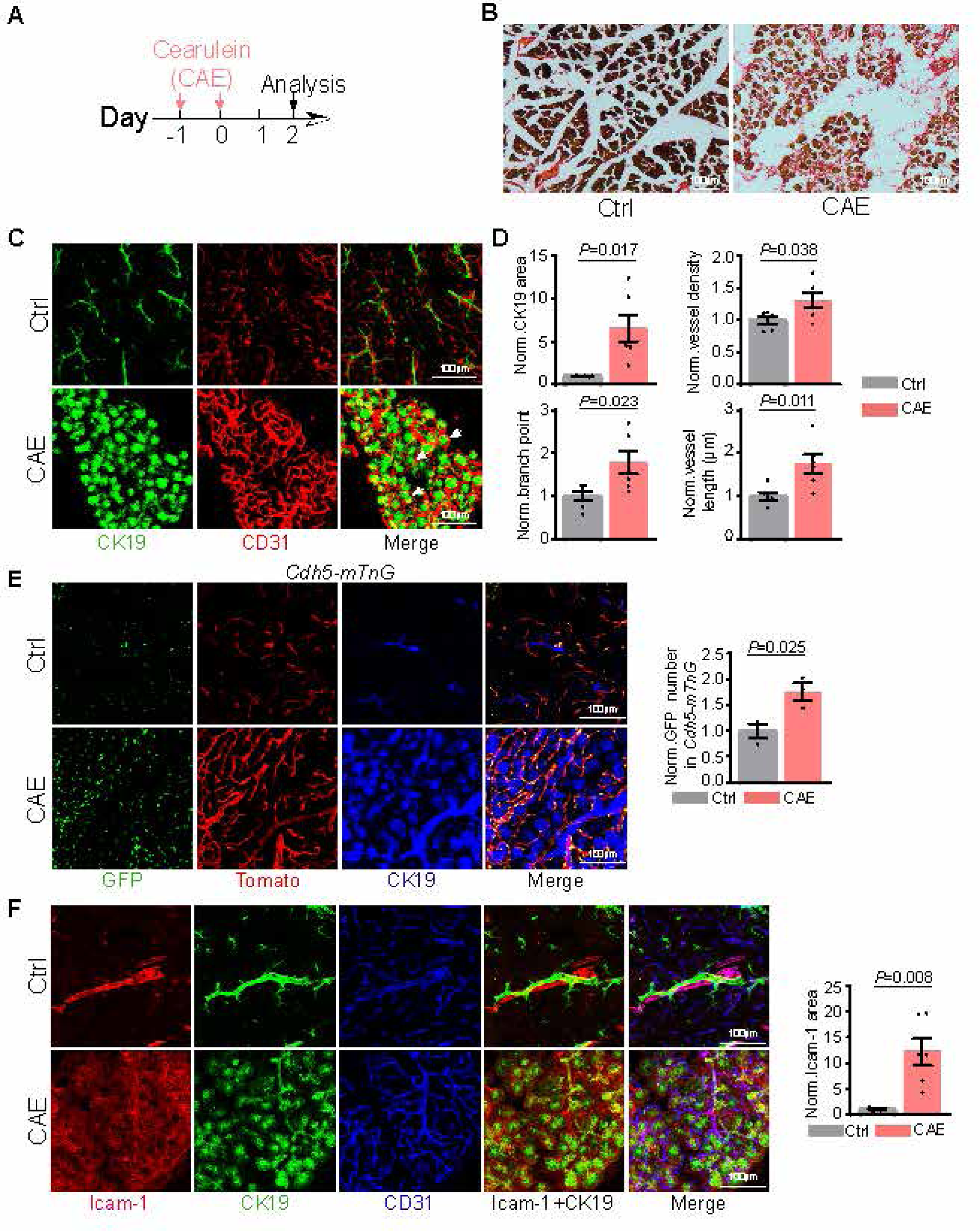
Cearulein-induced acute pancreatitis was associated with pathological angiogenesis. (**A**) Diagram depicting schedule for cearulein injection and analyzing time point. (**B**) Representative histological morphology of cearulein-treated (CAE) or vehicle treated (Ctrl) pancreas stained by sirius red. (**C**) Representative projected confocal images showing co-staining of CK19^+^ cells (green) together with CD31 (red) in cearulein-treated (CAE) or vehicle treated (Ctrl) pancreas. Arrows indicated the newly emergent CK19^+^ cells wrapped by intensive blood vessels. (**D**) Quantification about CK19^+^ covered area, length of blood vessels, number of vascular branch point and density of vasculature in cearulein-treated (CAE) or vehicle treated (Ctrl) pancreas. All data are normalized to ctrl. Ctrl=6; CAE=6. Error bars, mean ± s.e.m. *P* values, t-test. (**E**) Representative projected confocal images showing the number of endothelial cell nucleus visualized by nucleus-anchored GFP in cearulein-treated (CAE) or vehicle treated (Ctrl) pancreas of *Cdh5-mTnG* endothelial reporter mice. Green, nucleus GFP in *Cdh5-mTnG* mice; Red, tdTomato in *Cdh5-mTnG* mice; Blue, CK19. Quantification about GFP labelled endothelial cell number in cearulein-treated (CAE) or vehicle treated (Ctrl) pancreas of *Cdh5-mTnG* endothelial reporter mice. All data are normalized to ctrl. Ctrl=3; CAE=3. Error bars, mean ± s.e.m. *P* values, t-test. (**F**) Representative projected confocal images showing expression pattern of Icam-1 in cearulein-treated (CAE) or vehicle treated (Ctrl) pancreas. Red, Icam-1; Green, CK19; Blue, CD31. Quantification about Icam-1 covered area in cearulein-treated (CAE) or vehicle treated (Ctrl) pancreas. All data are normalized to ctrl. Ctrl=6; CAE=6. Error bars, mean ± s.e.m. P values, t-test.

In this model, we observed that the emerging CK19^+^ cells were wrapped by the vascular network, with a significant increase of CD31-labelled vascular length, vascular density and branch point numbers (Figure 2, C and D). A significant increase of VEC number was detected, with the help of *Cdh5-mTnG* reporter mice, in which the nucleus of VEC was genetically labelled with a GFP signal (Figure 2E). During acute pancreatitis, the area expressing Icam-1 was dramatically increased but the expression of Icam-1 seemed no longer restricted in pancreatic blood vessel (Figure 2F). The newly emergent CK19^+^ cells were surrounded by Icam-1^+^ cells during acute pancreatitis (Figure 2F). In contrast, the morphology and length of pancreatic artery did not show dramatic changes in response to acute pancreatitis (Supplemental Figure 2).

These evidences suggested that caerulein-induced acute pancreatitis was associated with pathological angiogenesis, which may represent an unexplored therapeutic target to prevent the progression of this disease.

### ScRNA-seq analysis revealed ductal-endothelial crosstalk as potential target to intervene pancreatitis progression

To investigate the potential of VEC as putative target for alleviating the symptom of acute pancreatitis, we utilized single cell RNA-sequencing (scRNA-seq) analysis to examine changes in the VEC transcriptomic profile as well as its communication with other pancreatic cells during the progression of acute pancreatitis(28). In the uniform manifold approximation and projection (UMAP) plot, acinar, ductal, islet and endothelial cells separated from each other with clear molecular markers to distinguish them (Figure 3A and Supplemental Figure 3, A to C). Molecular markers including CK19 (encoded by *Krt19*) were significantly upregulated in acinar cells on the first day after CAE treatment (CAE-D1) (Supplemental Figure 3D), which was consistent with the occurrence of ADM. Pearson’s correlation coefficient analysis showed that the overall cell variances between control and CAE treatment gradually decreased, consistent with the reversible feature of pancreatitis (Figure 3B).

**Figure 3.**
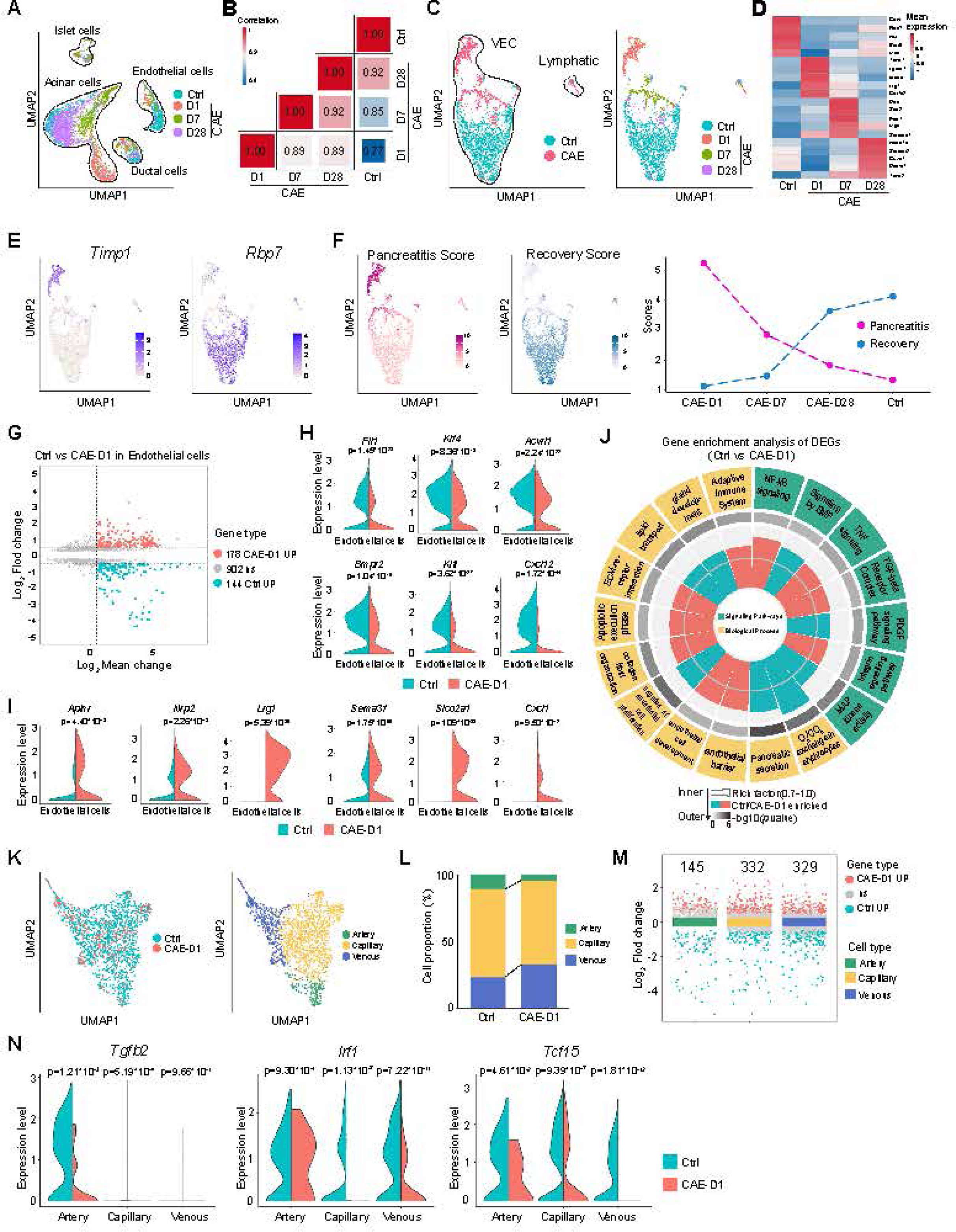
ScRNA-seq analysis of vascular endothelial cells in pancreas after cearulein treatment. (**A**) UMAP plot showing the dynamic changes of islet, ductal, acinar and endothelial cells in the ctrl or cearulein-treated pancreas at 1 day (CAE-D1), 7 days (CAE-D7), 28 days (CAE-D28) after treatment. (**B**) Pearson’s correlation coefficient analysis to compare the variance of all cells in different stages of pancreatitis with that in ctrl pancreas. (**C**) Visualization of the variance of endothelial cell in ctrl or CAE group in UMAP plot (left panel). UMAP plot showing the dynamic changes of endothelial cells in the ctrl or cearulein-treated pancreas at 1 day (CAE-D1), 7 days (CAE-D7), 28 days (CAE-D28) after treatment (right panel). (**D**) Heatmap showing the relatively enriched genes in the endothelial cell of ctrl, CAE-D1, CAE-D7, CAE-D28 pancreas. (**E**) UMAP plot showing *Timp1* and *Rbp7* as representative endothelial markers of CAE and Ctrl group, respectively. (**F**) UMAP plot showing combination of endothelial markers could evaluate the progression of pancreatitis. (**G**) MA plot showing the number of differentially-expressed genes (DEG) in VEC between ctrl and CAE-D1. (**H**) Violin plot showing the expression of *Fli1*, *Klf4*, *Acvrl1*, *Bmpr2*, *Kitl* and *Cxcl12* in VEC between ctrl and CAE-D1. (**I**) Violin plot showing the expression of *Aplnr*, *Nrp2*, *Lrg1*, *Sema3f*, *Slco2a1* and *Cxcl1* in VEC between ctrl and CAE-D1. (**J**) Gene ontology (GO) analysis of the biological process and signaling pathways enriched in ctrl or CAE-D1 VEC. (**K**) UMAP plot to compare the distribution of arterial, capillary and venous endothelial cell in ctrl and CAE endothelial cells. (**L**) Quantification of the percentage of each endothelial subpopulation in ctrl and CAE group. (**M**) The number of DEGs in different subpopulation of endothelial cells. (**N**) Violin plot showing the expression of *Tgfb2*, *Irf1* and *Tcf15* in different subpopulation of VEC between ctrl and CAE-D1.

Regarding VEC, the control and CAE-treated VEC were largely non-overlapping (Figure 3C and Supplemental Figure 3E), with molecular markers such as *Rbp7* and *Timp1* to distinguish them (Figure 3, D and E). Based on the combination of approximate 20 molecular markers, we could score the transcriptional feature of VEC to assess the degree of recovery after acute inflammation (Figure 3F). More than 300 differentially expressed genes (DEGs) were identified comparing control and CAE-treated VECs at day 1 (Figure 3G). At this stage, transcription factors including *Fli1*, *Id1*, *Klf4*, *Sox18*, and genes encoding crucial VEC receptors such as *Acvrl1*, *Bmpr2*, and *Bcam*, were significantly downregulated (Figure 3H and Supplemental Figure 3F). The expression of angiocrine factor, including *Kitl*, *Cxcl12*, and *Pdgfb*, were disrupted after CAE treatment (Figure 3H and Supplemental Figure 3F). Conversely, several genes promoting vascular growth including *Aplnr*(*17*), *Nrp2* (encoding Neuropilin 2)(29, 30), pathological angiogenesis marker *Lrg1*(*31*) and *Sema3f* (encoding Semaphorin 3f)(32, 33), were significantly upregulated (Figure 3I). The expression of organic anion transporter *Slco2a1* was dramatically increased after CAE treatment, suggesting the barrier function of VEC was changed (Figure 3I). Gene ontology (GO) analysis revealed an enrichment of genes related to pancreatic secretion function and lipid transport in control VECs, while genes associated with VEC development, function, and extracellular matrix interaction were enriched in CAE-D1 (Figure 3J). In control VECs, pathways like platelet-derived growth factor (PDGF), bone morphogenetic protein (BMP), and mitogen-activated protein kinase (MAPK) signaling were prominent, whereas integrin, tumor necrosis factor (TNF), and NF-κB signaling pathways were enriched in CAE-D1 (Figure 3J and Supplemental Figure 3G).

Sub-clustering analysis of pancreatic VEC divided them into arterial, venous and capillary clusters (Figure 3K and Supplemental Figure 3H). The proportion of these three VEC subpopulation was not dramatically altered in acute pancreatitis, with a slight increase of venous percentage (Figure 3L). DEG comparison suggested a stronger influence of CAE treatment on venous and capillary VECs than arterial VECs (Figure 3M), that was consistent with our immunostaining result (Figure 2 and Supplemental Figure 2). We noted that genes like *Tgfb2*, *Irf1*, and *Tcf15* were significantly reduced in arterial, capillary and venous VEC, respectively in CAE-D1 (Figure 3N), suggesting a subcluster-specific response to CAE treatment.

To investigate whether these transcriptomic changes affected the cell-cell communication in pancreas, we compared the putative crosstalk among acinar, endothelial, ductal and islet cells. Most prominently, CAE treatment dramatically reduced crosstalk numbers in endothelial, ductal, and islet cells (Figure 4A). Riverplot analysis about the details of these crosstalk highlighted that ductal and endothelial cells were major participants in cell-cell communication under both control and CAE conditions (Figure 4B). Notably, regardless of the condition, ductal cells were the primary interactors with VECs and vice versa (Figure 4C), that was consistent with the observed parallel alignment of ductal and endothelial cells in the pancreas (Figure 1). These evidences supported the existence of ductal-endothelial crosstalk in the pancreas. Next, we performed further analysis of ductal-endothelial crosstalk after caerulein treatment. The crosstalk from ductal to endothelial were dramatically reduced at 1 day after CAE treatment (Figure 4D). Within the crosstalk, certain putative signaling pathways, including junctional adhesion molecule-A, collagen-4, and macrophage migration inhibitory factor, maintained persistently, regardless of whether the pancreas was treated with CAE or not (Figure 4D). Notably, we detected a unique pathway involving the release of Spp-1 (or osteopontin) from ductal to endothelial cell targeting Integrin-α5. This signaling was selectively upregulated one day after CAE treatment, despite a dramatic overall reduction in crosstalk numbers, and it continued to be detected 28 days after CAE treatment (Figure 4, D and E). At endothelial subcluster level, CAE treatment reduced the number of cell-cell communication in artery, venous and capillary (Figure 4F). And the expression of Integrin-α5 was significantly increased in all three subpopulation of blood vessels after CAE treatment (Figure 4G).

**Figure 4.**
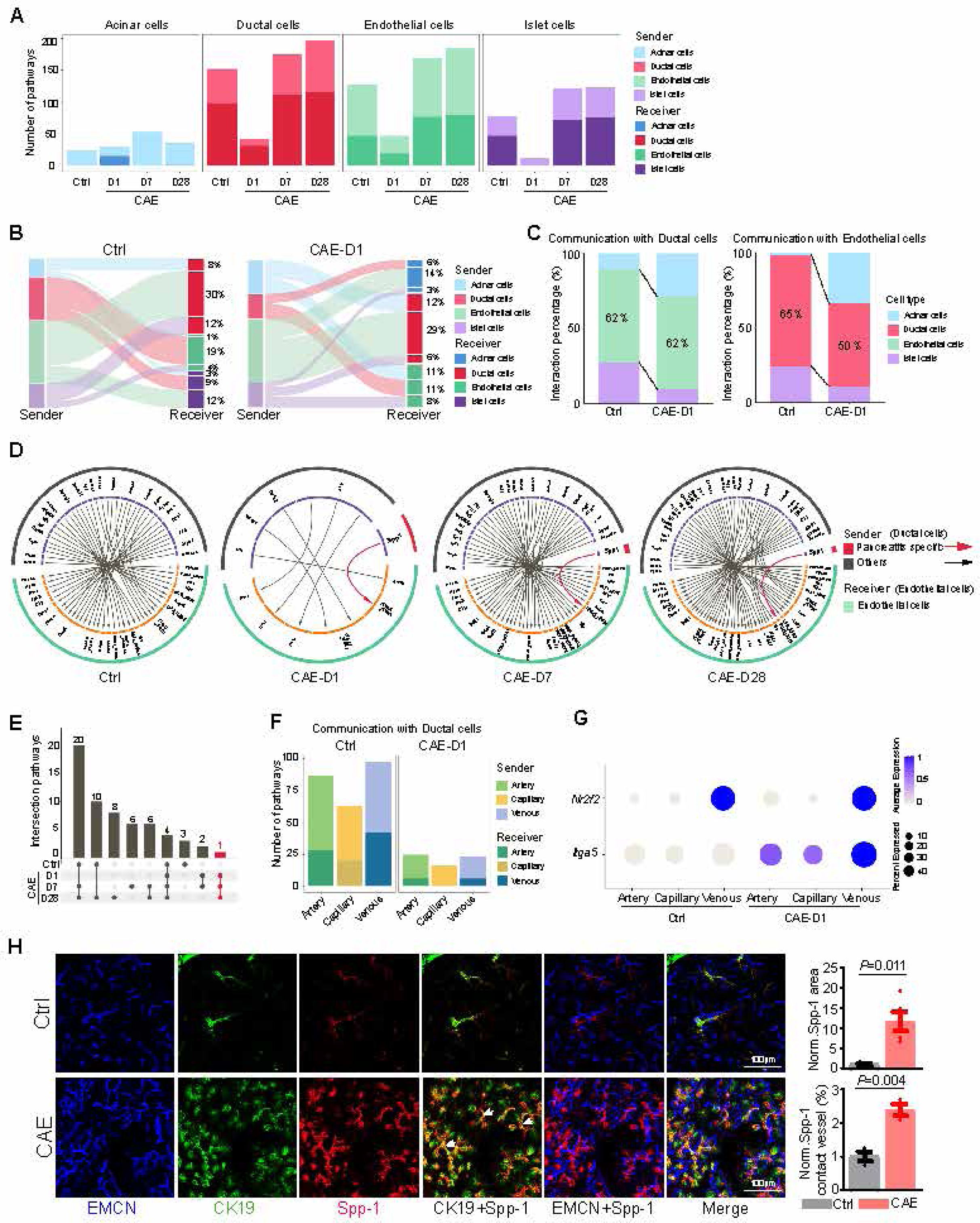
Cearulein treatment influence pancreatic cell-cell communication. (**A**) The number of putative crosstalk in acinar, ductal, endothelial and islet cells as signal sender or receiver. (**B**) River plot comparing the direction and percentage of putative crosstalk in acinar, endothelial, ductal and islet cells. (**C**) Quantitative comparison about the percentage of putative crosstalk in ductal cell and endothelial cell with other cell types between ctrl and CAE D1. (**D**) Visualization of putative crosstalk from ductal to endothelial cell in different stages of acute pancreatitis. Red line, pancreatitis enriched Spp-1/Integrin-α5 signaling. (**E**) Bar plot showing putative crosstalk from ductal to endothelial cells shared by different stages after cearulein treatment. Red bar, pancreatitis enriched Spp-1/Integrin-α5 signaling. (**F**) The number of putative crosstalk comparison between WT and CAE-D1 from ductal cell to different subpopulation of endothelial cells. (**G**) The expression of Integrin-α5 is significantly increased in different subpopulation of endothelial cells at 1 day after cearulein treatment. (**H**) Representative projected confocal images showing dramatic increase of Spp-1 release after cearulein treatment. Arrows indicated co-localization of CK19 and Spp-1. Quantification about Spp-1 covered area and Spp-1-contacted blood vessel length in cearulein-treated (CAE) or vehicle treated (Ctrl) pancreas. All data are normalized to ctrl. Ctrl=5; CAE=5. Error bars, mean ± s.e.m. P values, t-test.

To validate the finding from scRNA-seq analysis, we conducted immunostaining and found that there was a significant increase in the expression level of Spp-1, the Spp-1 positive area, as well as the length of blood vessels in contact with Spp-1 after CAE treatment (Figure 4H). The Spp-1 signals were predominantly co-localized with CK19 (Figure 4H), supporting our results that ductal or ductal-like CK19^+^ cells were the major source of this paracrine factor in the pancreas.

Taken together, the scRNA-seq analysis revealed that the transcriptomic profile of pancreatic VEC were significantly changed in CAE-induced acute pancreatitis. In the pancreas, the interplay between ductal and endothelial cells was one of the most prevalent forms of cell-cell communication, which was disrupted during acute pancreatitis, with the exception of the Spp-1/Integrin-α5 signaling pathway.

### ATN-161 mitigates the pathological defects in caerulein-induced pancreatitis

To investigate the importance of ductal-endothelial crosstalk, particularly the Spp-1/Integrin-α5 signaling during the progression of acute pancreatitis, we performed the following experiments using ATN-161, a specific antagonist of Integrin-α5(34), to block the Spp-1/Integrin-α5 signaling. This compound was intravenously administered concurrently with CAE injections (Figure 5A). After ATN-161 treatment, we detected a significant normalization of pancreatic morphology, which included a notable reduction in red collagen fibers and a return to more polygonal tissue morphology, reminiscence of the normal pancreatic tissue staining observed by Sirius red staining (Figure 5B and 2B). Remarkably, the number of CK19^+^ pancreatic cells decreased significantly following ATN-161 treatment in the pancreatitis model, suggesting that ADM was inhibited by this antagonist (Figure 5, C and D). Together with the reduction in CK19^+^ cell numbers, other pathological phenomena induced by CAE treatment, such as vascular density, vascular length, Icam-1^+^ area, Spp-1 release, and Spp-1-contacted vessels, were all significantly reduced (Figure 5, C to F), suggesting the symptoms of acute pancreatitis was substantially mitigated. To confirm the influence of Spp-1 and ATN-161 on VEC, we treated human umbilical vein endothelial cells (HUVEC) with these two chemicals. Spp-1 treatment promoted wound healing ability of HUVEC, but this promotion was largely abolished when ATN-161 was added to the medium (Figure 5, G and H), suggesting a direct influence of Spp-1 and ATN-161 on VEC.

**Figure 5.**
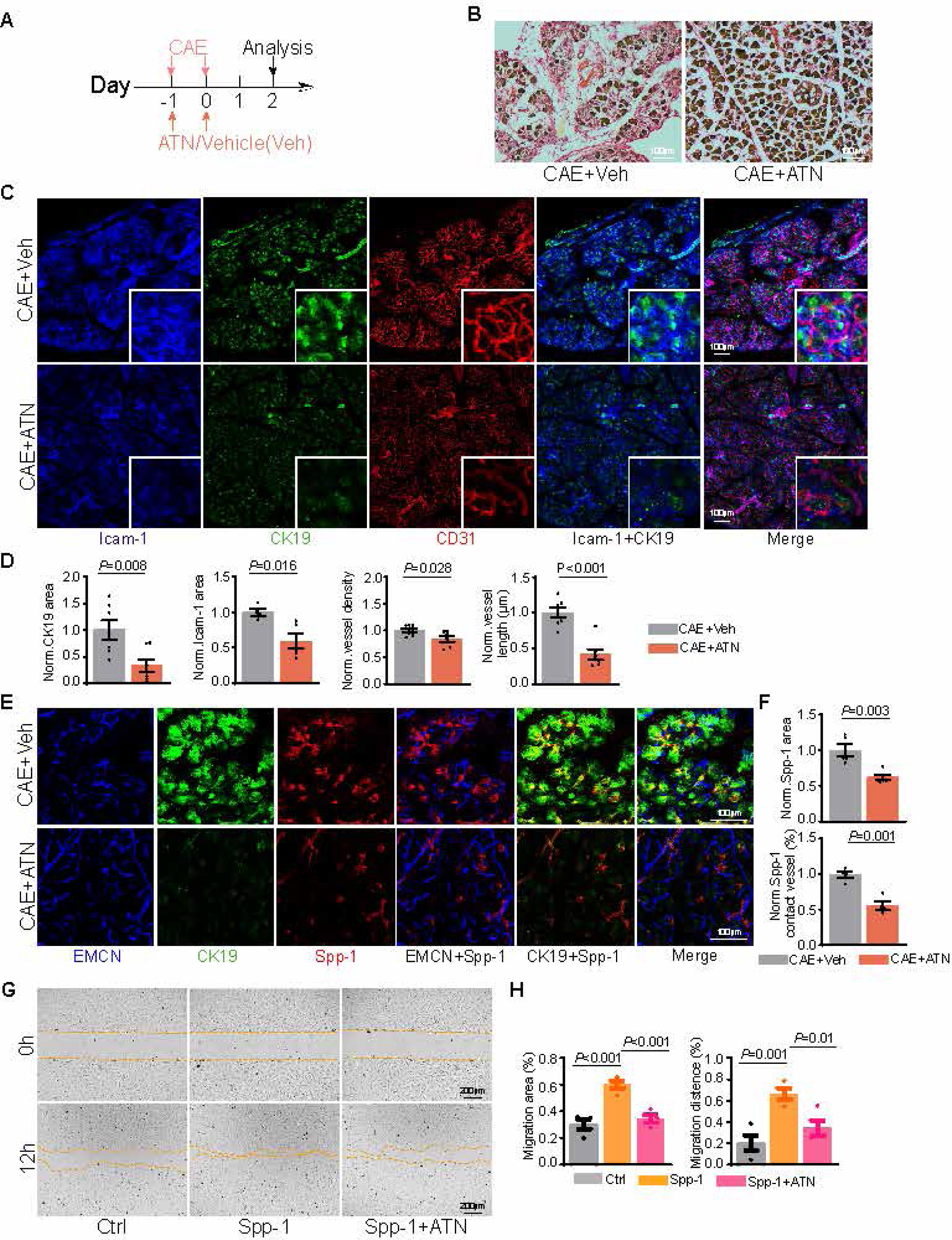
ATN-161 treatment alleviates cearulein-induced pancreatic disorder. (**A**) Diagram depicting schedule for co-injection of cearulein and ATN-161. (**B**) Representative histological morphology of vehicle and ATN-161-treated pancreas stained by sirius red in cearulein model. (**C**) Representative projected confocal images showing expression of Icam-1 (blue), CK19 (green) and CD31 (red) in vehicle and ATN-161-treated pancreas of cearulein model. The image in square showing figures with higher magnification. (**D**) Quantification about CK19^+^ covered area, length of blood vessels, vascular density and CK19^+^ covered area in vehicle and ATN-161-treated pancreas of cearulein model. All data are normalized to ctrl. CAE+Vehicle=7; CAE+ATN=7. Error bars, mean ± s.e.m. P values, t-test. (**E**) Representative projected confocal images showing expression of CK19 (green) and Spp-1(red) in vehicle and ATN-161-treated pancreas of cearulein model. (**F**) Quantification about Spp-1^+^ covered area and length of Spp-1-contacted vessel in vehicle and ATN-161-treated pancreas of cearulein model. All data are normalized to ctrl. CAE+ Vehicle =5; CAE+ATN=5. Error bars, mean ± s.e.m. P values, t-test. (**G**) Representative images showing the wound healing in HUVEC when Spp-1 or Spp-1 together with ATN-161 was supplement to the culture medium. (**H**) Quantification about the migration area and migration distance in ctrl (N=4), Spp-1 (N=4), Spp-1+ATN-161 (N=4) treatment group.

To objectively evaluate the situation of pancreas after ATN-161 treatment in the pancreatitis model, we performed scRNA-seq analysis to assess the transcriptomic status of these pancreatic cells. In the integrated UMAP plot, most cells co-injected with CAE and ATN-161 (abbreviated as ATN group) did not overlap with cells at CAE-D1 (Figure 6A and Supplemental Figure 4, A and B). Pearson’s correlation coefficient analysis revealed that the ATN group was more similar to the overall transcriptomic status at CAE-D28 with a similarity as high as 0.95 (Figure 6B). Consistent with the immunostaining result (Figure 5C), CK19 (*Krt19*) expression was significantly reduced in the ATN group (Supplemental Figure 4C). In UMAP plot, the VECs from ATN and CAE-D1 were clearly separated from each other (Figure 6C). Scoring evaluation indicated the status of VEC in the ATN group was between CAE-7D and CAE-28D (Figure 6D and Supplemental Figure 4, D and E). Over 400 differentially-expressed genes were identified in VEC (Figure 6E and Supplemental Figure 4, F and G), with notable downregulation of *Flt1* (encoding VEGFR1), *Kdr* (encoding VEGFR2), suggesting the efficacy of ATN-161 in inhibiting pancreatic pathological angiogenesis (Figure 6F). ATN-161 also attenuated the expression of other angiogenesis-related genes including *Tie1*, *Eng*, *S1pr1*, *Plvap*, *Klf2*, *Atf3* and *Fzd4* (Figure 6F and Supplemental Figure 4H). While ATN-161 blocked the transcription of angiocrine factors like *Cxcl1* and *Tgfb1* (Figure 6F), other angiocrine factors such as *Ang*, *Ptn*, and *Ccl11* were significantly upregulated (Figure 6G). Transcription factor Pbx1, carboxypeptidase and gene (*Mfap5*) associated with VEC adhesion function(35) were dramatically enhanced in the VEC of ATN group (Figure 6G). After ATN-161 treatment, biological processes related to pancreatic development, pancreatic secretion function, and cell proliferation were rescued, whereas pathways related to vascular development were relatively higher in CAE-1D comparing to ATN group (Figure 6H and Supplemental Figure 4, I and J). Pathways including Apelin, Integrin, and NF-κB signaling were more prominent in CAE-1D (Figure 6H). Notably, transcriptions of *Id1*, *Id3*, *Sox18*, *Fli1* and *Fbln2*, significantly downregulated by CAE treatment, were apparently restored after ATN-161 co-injection (Figure 6I). Meanwhile, CAE treatment significantly promoted the expression of *Aplnr*, *Flt4*, *Ephb4*, *Itga5*, *Icam1* and *Cldn5* but ATN-161 normalized the expression of these genes in pancreatic VEC (Figure 6J).

Regarding the crosstalk, ATN-161 treatment notably restored the reduction in cell-cell communication (Figure 6, K and L). The ductal-endothelial crosstalk occupied 60% of cell-cell communication in ATN group (Figure 6M). Although acinar cells still functioned as signal receiver in ATN group, its percentage had reduced from 23% to 13% after ATN-161 treatment (Figure 4B and 6M). The islet cells had recovered their signal receiver function in ATN group (Figure 4B and 6L). The percentage of signaling from ductal to endothelial cell had recovered from 37% to 49% after ATN-161 treatment (Figure 6N). In the crosstalk, we noted that 17 pathways were shared by control, CAE-7D, CAE-28D with ATN group (Figure 6, O and P). These evidences suggested that the cell-cell communication in pancreas had largely recovered after ATN-161 treatment. More specifically, the signal strength of laminin-α4, collagen and class-3 semaphorin signaling had recovered while the increase of Spp-1/Integrin-α5 signaling had been significantly blocked by ATN-161 treatment (Figure 6Q).

**Figure 6.**
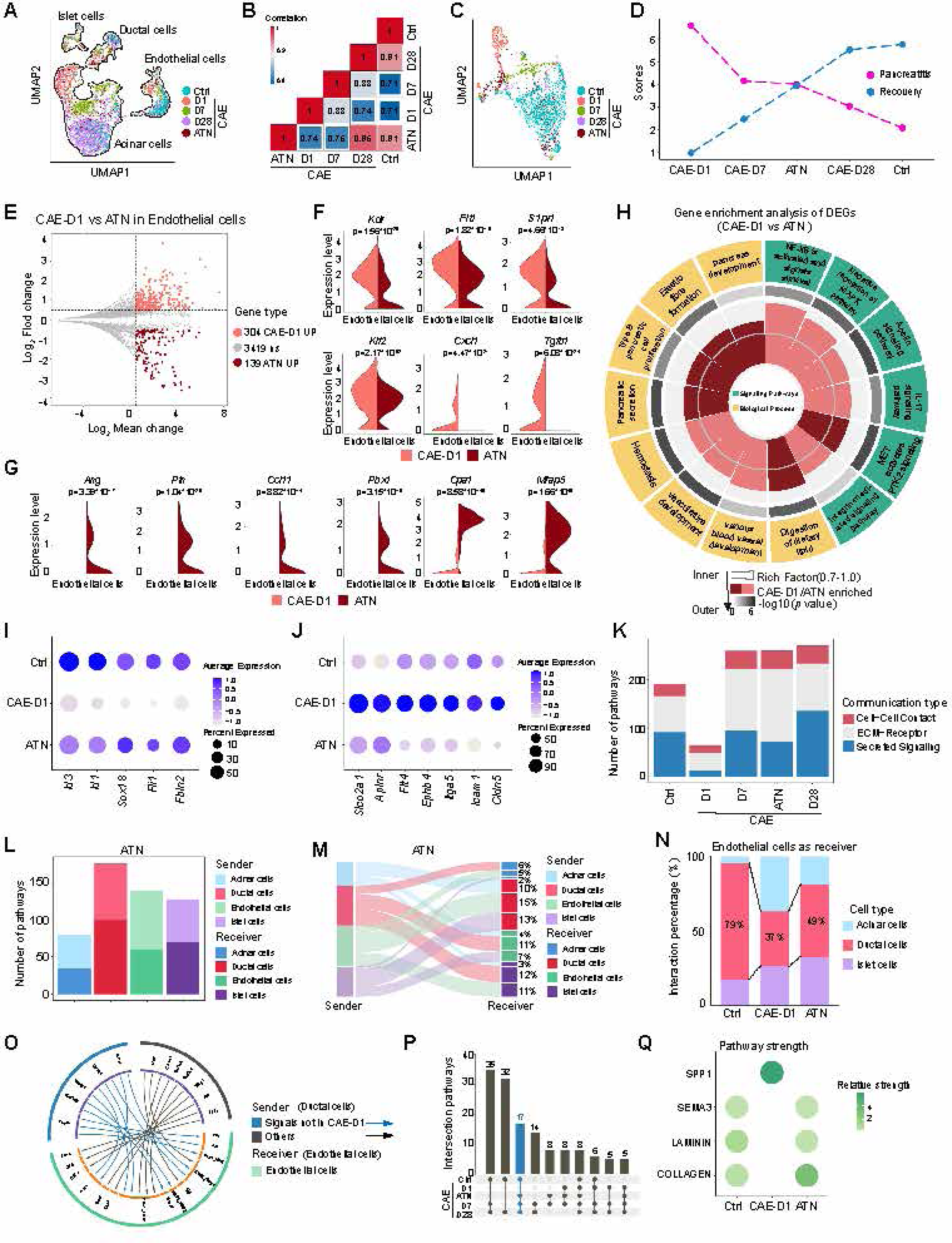
ScRNA-seq analysis of pancreas after ATN-161 treatment in cearulein-induced acute pancreatitis. (**A**) UMAP plot showing the dynamic changes of islet, ductal, acinar and endothelial cells in the ctrl, cearulein-treated pancreas at 1 day (CAE-D1), 7 days (CAE-D7), 28 days (CAE-D28) after treatment or cearulein with ATN-161 co-injected pancreas at 2 days after treatment. (**B**) Pearson’s correlation coefficient analysis to compare the variance of all cells in different condition of pancreatitis with that in ctrl pancreas. Note that ATN group has highest similarity with CAE-D28 cells. (**C**) UMAP plot showing the distribution of CAE+ATN treated VEC together with the dynamic changes of VEC in the ctrl or CAE-treated pancreas at 1 day (CAE-D1), 7 days (CAE-D7), 28 days (CAE-D28) after treatment. (**D**) Scoring to evaluate the transcriptomic status of ATN-treated VEC. Both the pancreatitis and recovery scores fall between CAE-D7 and CAE-D28. (**E**) MA plot showing the number of differentially-expressed genes (DEG) in VEC between CAE-D1 and ATN. (**F**) Violin plot showing the expression of *Kdr*, *Flt1, S1pr1, Klf2, Cxcl1* and *Tgfb1* in VEC between CAE-D1 and ATN. (**G**) Violin plot showing the expression of *Ang, Ptn, Ccl11, Pbx1, Cpa1* and *Mfap5* in VEC between CAE-D1 and ATN. (**H**) Gene ontology (GO) analysis of the biological process and signaling pathways enriched in CAE-D1 and ATN. (**I**) Dot plot showing ATN-161 treatment restores the expression of *Id1, Id3, Sox18, Fli1* and *Fbln2*. (**J**) Dot plot showing ATN-161 treatment normalizes the expression of *Slco2a1, Aplnr, Flt4, Ephb4, Itga5, Icam1* and *Cldn5*. (**K**) The number of putative crosstalk in acinar, endothelial, ductal and islet cells in the ctrl, CAE-D1, CAE-D7, CAE-D28 and ATN groups. Note that the number of crosstalk falls between CAE-D7 and CAE-D28. (**L**) The number of putative crosstalk in acinar, endothelial, ductal and islet cells as signal sender or receiver in ATN group. (**M**) River plot comparing the direction and percentage of putative crosstalk in acinar, endothelial, ductal and islet cells in ATN group. (**N**) Quantitative comparison showing ATN-161 restores the percentage of putative crosstalk sending from ductal cells and to endothelial cells. (**O**) Visualization of putative crosstalk from ductal to endothelial cell in ATN group. (**P**) Bar plot showing putative crosstalk from ductal to endothelial cells shared by different stages of CAE treatment. Blue bar shows the number of shared pathways by all the other stages except CAE-D1. (**Q**) Scoring the pathway strength showing that ATN-161 treatment restores semaphorin, laminin and collagen signaling and normalize the Spp-1 signaling.

To sum up, these evidences suggested that ATN-161 had a robust effect to alleviate the pathological characteristics during caerulein-induced acute pancreatitis.

### Yohimbine-induced extension of pancreatitis is disrupted by ATN-161

Although caerulein administration is a well-established model for acute pancreatitis, its effects are reversible, limiting the ability of this model to evaluate the impact of ATN-161 in more prolonged pancreatitis conditions.

To address this question, we found that intraperitoneally injection of caerulein together with yohimbine (abbreviated as YOH), a potent and alpha 2-adrenergic receptor antagonist(36), extended the symptom of pancreatitis (Figure 7A). In the vehicle control, the number of CK19^+^ pancreatic cells already decreased from the peak at 5 days after final caerulein injection (Figure 7B). However, when caerulein was co-injected with yohimbine, there was a sustained increase in CK19^+^ cells (Figure 7B). Notably, the morphology of these CK19^+^ cells changed from circle to tubular structures (Figure 7B and 2C), suggesting yohimbine co-injection induced an essential progression of ADM. Other pathological phenotypes, such as vascular density, Icam-1^+^ area, Spp-1^+^ area, and CK19^+^ area, were similarly persistent in the yohimbine-induced extended pancreatitis model (Figure 7, B to E).

**Figure 7.**
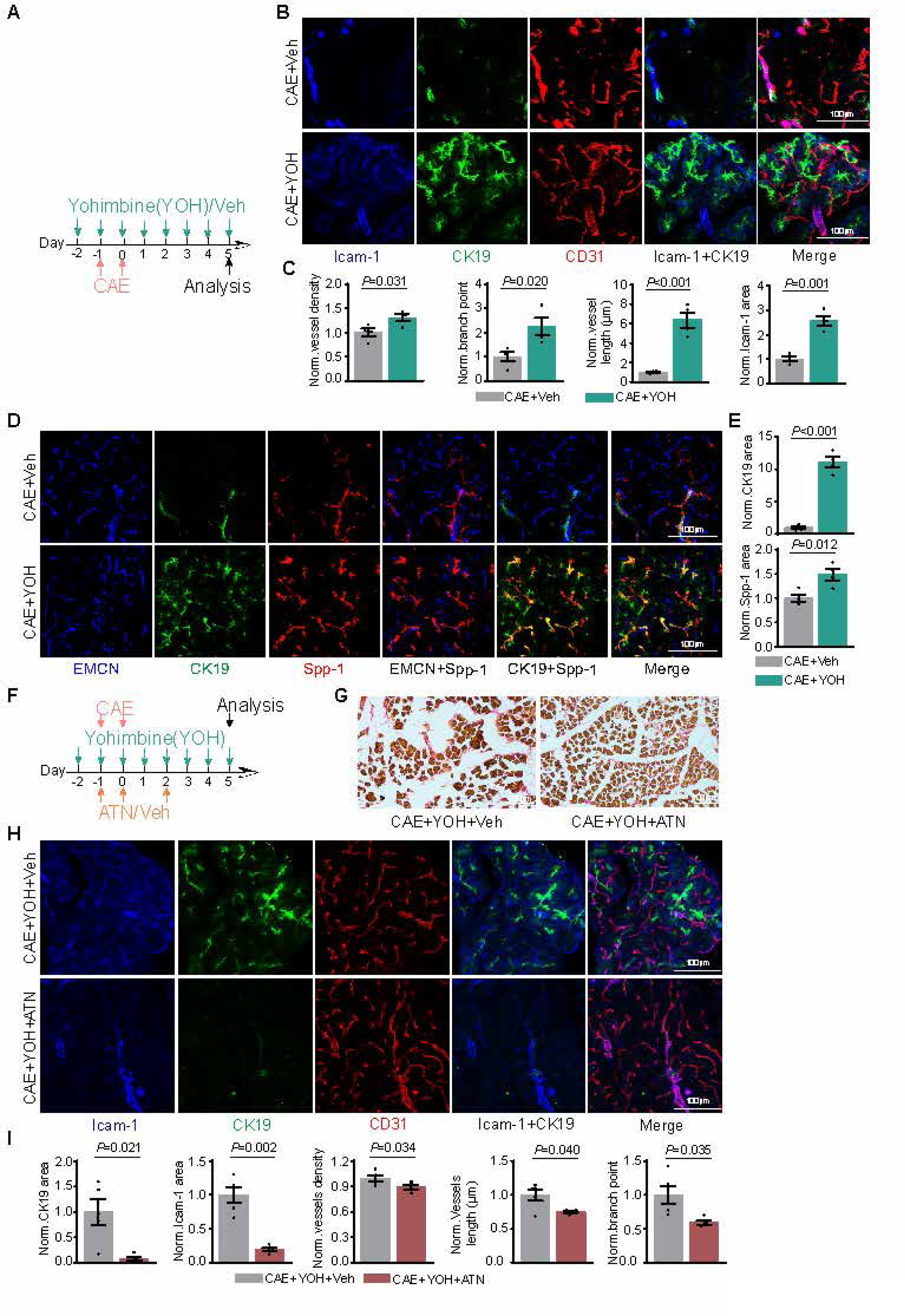
ATN-161 mitigates yohimbine and cearulein induced prolonged-pancreatic disorder. (**A**) Diagram depicting the protocol to inject cearulein and yohimbine. (**B**) Representative projected confocal images showing expression of Icam-1 (Blue), CK19 (Green) and CD31(Red) in vehicle and yohimbine-treated pancreas of cearulein model. Yohimbine, YOH. (**C**) Quantification about vascular length, vascular density, vascular branch point number and Icam-1^+^ covered area in vehicle and yohimbine-treated pancreas of cearulein model. All data are normalized to ctrl. Yohimbine, YOH. CAE+DMSO=4; CAE+YOH=4. Error bars, mean ± s.e.m. *P* values, t-test. (**D**) Representative projected confocal images showing the expression of EMCN (Blue), CK19 (Green) and Spp-1 (Red) in vehicle and yohimbine-treated pancreas of cearulein model. Yohimbine, YOH. (**E**) Quantification about CK19^+^ and Spp-1^+^ area in vehicle and yohimbine-treated pancreas of cearulein model. All data are normalized to ctrl. Yohimbine, YOH. CAE+DMSO=4; CAE+YOH=4. Error bars, mean ± s.e.m. *P* values, t-test. (**F**) Diagram depicting the protocol to inject cearulein, ATN-161 and yohimbine. (**G**) Representative histological morphology of vehicle and ATN-161 in yohimbine-induced prolonged pancreatitis. Sample were stained by sirius red. (**H**) Representative projected confocal images showing the expression of Icam-1 (blue), CK19 (green) and CD31 (red) in vehicle and yohimbine-treated pancreas of cearulein model. Yohimbine, YOH. (**I**) Quantification about Icam-1^+^ area, CK19^+^ area, vascular density, vascular length, vascular branch point number in vehicle and ATN-161-treated yohimbine-induced prolonged pancreatitis. All data are normalized to ctrl. Yohimbine, YOH. ATN-161, ATN. CAE+YOH+PBS=5; CAE+YOH+ATN=5. Error bars, mean ± s.e.m. *P* values, t-test.

To evaluate the efficacy of ATN-161 in this extended pancreatitis model, we co-injected ATN-161 together with caerulein and yohimbine (Figure 7F). Morphologically, ATN-161 treatment led to an increase in the polygonal tissue area in the pancreas (Figure 7G). Similarly, the number of CK19^+^ cells, vascular density, vascular length and Icam-1^+^ area were all reduced when ATN-161 was administrated in conjunction with caerulein and yohimbine (Figure 7, H and I).

These findings suggest that yohimbine treatment enhances the tubular morphogenesis of CK19^+^ cells and prolongs the pathological condition of pancreatitis. ATN-161 is effective to mitigate the extended symptoms of pancreatitis, reinforcing its potential as a therapeutic agent in more extended case of pancreatitis.

## Discussion

As one of the most prevalent gastrointestinal disorders, the global burden of pancreatitis is very severe, with approximate 7 to 9 million patient leading to billions of US dollar’s health care cost as well as 40% mortality in severe cases(5–9). However, pharmacological interventions to prevent the progression of pancreatitis are very limited.

In this work, we observe that CK19^+^ ductal epithelial cells align parallel to blood vessels in the pancreas. This spatial association facilitates intense crosstalk between ductal and endothelial cells, as supported by our scRNA-seq cell-cell communication analysis. During acute pancreatitis, the emergence of CK19^+^ cells through acinar-to-ductal metaplasia (ADM) is coupled with pathological angiogenesis. This process is triggered, at least in part, by the release of Spp-1 from CK19^+^ cells, activating Integrin-α5 signaling in pancreatic VECs. The blockade of this signaling pathway by ATN-161, a specific antagonist of Integrin-α5, significantly restrains ADM and pathological angiogenesis. Consequently, this restores vascular homeostasis, pathological gene expression, and cell-cell communication in acute pancreatitis model. Finally, our study reveals that the potent alpha-2 adrenergic receptor antagonist yohimbine can extend the symptoms of caerulein-induced pancreatitis and promote tubular morphogenesis of CK19^+^ cells. This extended pathological condition can also be mitigated by ATN-161. Our work reveals the significance of crosstalk between ductal and endothelial cells in pancreatitis, which could be targeted by ATN-161 as pharmacological intervention approach to alleviate pancreatitis.

In the last two decades, numerous studies have reported the parallel alignment of blood vessel and peripheral nerve, forming a neurovascular interface that facilitates their communication and tissue morphogenesis(22, 37, 38). However, as far as we know, reports of other cell types running parallel to blood vessels are scarce. Our work demonstrates that pancreatic ducts, composed of ductal epithelial cells and responsible for transporting pancreatic juice, form a previously unrecognized network sharing spatial association with blood vessels.

Previous works have shown associations between blood vessels and pancreatitis, such as ischemia or blood reperfusion leading to pancreatitis(39). Pancreatitis could induce splenic vein thrombosis(40) or life-threatening hemorrhage(41, 42). However, the cell-cell communication between VECs and pancreatic resident cells remain largely unexplored. In other organs, such as bone marrow, brain and liver, VEC release several angiocrine factors to target the tissue resident cells to mediate organ development and regeneration(43–47). Hematopoietic stem/progenitor cells, astrocyte and cardiomyocyte derived signals are reported to induce differentiation of VEC and angiogenesis(17, 48, 49). Our findings suggest that ductal and endothelial cells in the pancreas form an intimate partnership for crosstalk, which could be targeted to ameliorate the progression of pancreatic diseases. Since pancreatic ductal cells are involved in various diseases including pancreatitis and pancreatic ductal adenocarcinoma, targeting the ductal-endothelial interface may offer more therapeutic options.

Secreted phosphoprotein 1 (Spp-1), namely osteopontin, is initially identified as one of the noncollagenous proteins present in bone matrix(50). Recent researches have significantly expanded our knowledge of this secretion protein in cardiovascular disease, cancer and diabetes(50–52). Spp-1 levels are known to increase in the serum or plasma of human patient with acute pancreatitis and pancreatic cancer (53–56), which is consistent with our result (Figure 4H). However, the source and downstream target of Spp-1, especially whether this signaling pathways could be intervened to alleviate the progression of pancreatic disease has not been thoroughly investigated. Integrins, a large family of receptors, bind various extracellular matrix and cell surface proteins. Spp-1 can directly bind and activate the downstream target of several integrins including Integrin-α5 on several different types of cells(57–61). In this work, we found that Spp-1/Integrin-α5 signaling was significantly strengthened during acute pancreatitis (Figure 4).

ATN-161, a five–amino-acid peptide that preferentially inhibit Integrin-α5, has been utilized as antiangiogenic peptide(58, 62). ATN-161 is well tolerated by human patient in the phase 1 clinical trial, in which nine patients received more than 100 days of ATN-161 treatment for thrice every week (63), which warrant the tolerability and safety of ATN-161. Although ATN-161 has shown limited effect on solid tumors, our research indicates its efficacy in mitigating the progression of acute pancreatitis, suggesting its potential as a therapeutic option to prevent the escalation of severe pancreatitis.

In summary, our study demonstrates the spatial association and signaling crosstalk between ductal and vascular endothelial cells in the pancreas. Targeting the Spp-1/Integrin-α5 signaling pathway with ATN-161 emerges as a promising therapeutic strategy to alleviate the symptoms of pancreatitis.

## Methods

### Animal experiments

*Cdh5(PAC)-mtdTomato-nGFP* (*Cdh5-mTnG*) endothelial cell reporter mice were previously used to label endothelial cell with membrane tdTomato signal and endothelial cell nucleus with nucleus GFP signal(17). *Bmx-CreERT2*(64) transgenic mice were interbred with the *Rosa26-mTmG*(65) strain to generate *Bmx-CreERT2 Rosa26-mTmG* for genetic labelling of arterial ECs. 1mg tamoxifen were administered on 5 consecutive days to induce GFP expression. Genetically modified animals were genotyped using respective PCR protocols and were available on reasonable request.

8-12 week-old male C57BL/6 mice were used as wild-type young adults. 75 µg/kg caerulein were administrated for 16 times on 2 consecutive days to induce acute pancreatitis. 10 mg/kg yohimbine were administrated on 8 consecutive days to induce prolonged acute pancreatitis. Approximate 8 mg/kg ATN-161 were injected on 2 consecutive days to wild-type mice. These drugs were administrated following the schedules illustrated in corresponding figures.

Animals were housed in the animal facility of Guangzhou Institutes of Biomedicine and Health (GIBH). Animal experiments were performed according to the institutional guidelines and laws, following the protocols (2023075) approved by local animal ethics committees.

### Cryosectioning, immunostaining and confocal imaging

Pancreas were dissected and placed in ice-cold 4% paraformaldehyde (PFA-PBS) (Sigma, P6148) solution and fixed overnight. Fixed pancreas samples were dehydrated in 20% sucrose-PBS solution, and embedded in OCT (SAKURA, 4583) for storage at −80°C. Cryosectioning was performed on a Leica CM3050 cryostat with low profile blades to cut 40 µm. For immunostaining, pancreatic sections were rehydrated in PBS, permeabilized for about 15 min in 0.5% Triton X100-PBS solution and blocked for about 2 hours in PBS with 1% BSA, 2% donkey serum, 10% Trion X100-PBS (blocking buffer) at room temperature. Sections were stained with primary antibodies diluted in blocking buffer at 4°C overnight. After incubation, sections were washed three times with PBS and incubated with appropriate secondary antibodies diluted in blocking buffer at room temperature for 2 hours. Nuclei were stained with DAPI during secondary antibody incubation. After that, sections were washed three times with PBS, mounted with Fluoromount-G (0100-01, Southern Biotech) and kept in 4°C for confocal imaging. The primary antibodies include Emcn (Santa Cruz, sc-65495, 1:100), CD31 (R&D, AF3628, 1:100), VEGFR2 (R&D, AF644, 1:60), Spp-1 (R&D, AF808, 1:150), Icam-1 Antibody (Biolegend, 116101, 1:60), Anti-Cytokeratin19 (Abcam, AB52625, 1:80) were used in this study. The secondary antibodies include Donkey anti-Rat Alexa Fluor 488 (Thermofischer Scientific, A21208, 1:200), Donkey anti-Rabbit Alexa Fluor 546 (Thermofischer Scientific, A10040, 1:200), Donkey anti-Goat Alexa Fluor 647 (Thermofischer Scientific, A21447, 1:200), Donkey anti-rabbit Alexa Fluor 647 (Thermofischer Scientific, A31573, 1:200), Donkey anti-Rat IgG (H+L) Highly Cross-Adsorbed Secondary Antibody, Alexa Fluor™ 594 (Thermofischer Scientific, A21209, 1:200). Sections were imaged with laser scanning confocal microscopes (LSM710, LSM800) after immunohistochemistry. Quantitative analysis of drug-treated phenotypes was imaged with the same microscope and imaging acquisition setting for treatment and control samples.

We used Fiji (open source; http://fiji.sc/) and AngioTool (https://angiotool.software.informer.com/) for image processing in compliance with guide for digital images. In general, three technical replicate areas were randomly chosen from one biological sample for analysis. Quantification of Icam-1, CK19 and Spp-1 stained area was performed in Fiji. Images exported from Fiji were used to quantify vascular density, vascular length, branch point in AngioTool.

### scRNA-seq sample preparation

Pancreases were dissected and cut by knife before cells were collected in 10% FCS-PBS solution with protease inhibitor cocktail (Beyotime, P1005). The tissue was immersed in dissociation solution (5% FCS-PBS solution with collagenase Ⅳ, collagenase Ⅰ, DNase Ⅰ, dispase Ⅱ) and incubated at 37°C for 30 minutes. The dissociated cells were collected into 10% FCS-PBS solution. Samples were filtered using 100 μm Nylon cell strainer (Biosharp, BS-100-XBS) to get single cell suspensions, and cell centrifugation was started at 300 g for 5 min, gradually decreasing the speed of centrifugation. Subsequently, the primary antibody include CD45-PE/Cy7 (Biolegend, 103114), Ter119-PE/Cy7 (BD pharmingen, 557853), APC anti-mouse CD31 Antibody (Biolegend, 102410), PE/Cyanine7 anti-mouse CD140a Antibody (Biolegend, 135911) were diluted with 5% FCS-PBS solution and incubated with the cells on ice for 20 min, then the excess primary antibody was washed away. AO/PI was added to check cell viability. Afterwards, live cells and CD45^-^TER119^-^ CD31^+^ cells were sorted on an BD FACS Arial Sorter.

### ScRNA sequencing analysis

Sequencing data for ATN group were aligned to the mouse reference genome (mm10) and quantified by the STAR software package (version 2.7.10b) with default parameters. After the genome alignment, we recovered 42291 reads per cell. The sequencing saturation percentage was 95%.

Data normalization, detailed analysis, and visualization were performed using the Seurat package (version 4.3.0)(66) unless otherwise specified. For the initial quality control of the extracted gene-cell matrices, cells were filtered based on parameters nFeature_RNA > 200 & nFeature_RNA < 5000 for the number of genes per cell, and percent.mt < 10 for the percentage of mitochondrial genes. Additionally, genes with parameters min.cell = 3 and min.features = 200 were considered. The filtered matrices were normalized using the LogNormalize method with a scale factor of 10000. Variable genes were identified with default options and used for principal component analysis. The first 7 principal components were used for UMAP non-linear dimensional reduction. FindIntegrationAnchors and IntegratedData with default options were used for data integration. Unsupervised hierarchical clustering analysis was performed using the FindClusters function in the Seurat package. The cellular identity of each cluster was determined by finding cluster-specific marker genes using the FindAllMarkers function and comparing those markers to known cell type-specific genes from previous studies. The mean of different cluster-specific genes was visualized using the pheatmap (version 1.0.12) package. FeaturePlot, VlnPlot, and DotPlot functions of the Seurat package were used for the visualization of selected genes.

To detect differentially expressed genes (DEGs) in different clusters of datasets, the FindAllMarkers function was used. Genes with an avg_logFC (log fold-change in the average expression) > 0.5 and log2mean (log mean of gene expression) > 0.5 were considered as DEGs. Gene ontology analysis was performed using Kobas (version 3.0)(67) software; we selected p-value < 0.05 for visualization. The top 10 DEGs of endothelial cells between Ctrl and CAE-D1 were selected using the AddModuleScore function of Seurat to perform the pancreatitis and recovery feature scores. These scores were then rescaled to a 0-10 scale across all sample cells. Col1a1, Col1a2, Rbp1, Btnl9, Car4, and Rbp7 were scored, and the mean of each gene in each condition was calculated and visualized using the ggplot2 (version 3.4.0) package.

Pearson correlation analysis using differentially expressed genes between groups was conducted to identify the similarity of different conditions. First, the mean of each gene within each group was calculated. Second, the top 2000 DEGs between different groups were selected. Third, the cor function with default options in R was used to calculate the correlation of groups. Finally, the results were visualized using the pheatmap package.

The CellChat (version 1.6.1)(68) software was employed to predict ligand-receptor interplays and cellular communication networks. Default parameters were used during the cell-cell interaction analysis. Interactions across cell types in each condition were shown in a bar plot and river plot using the ggplot2 package. The ligand-receptor plot was visualized using the circlize (version 0.4.15) package.

### Cell Culture

The human umbilical vein endothelial cells (HUVEC) were purchased from ATCC, cultured in ECM medium (ScienCell,1001) with extra 10% Fetal Bovine Serum (FBS) (NEWZERUM, FBS-E500) and grown in 5% CO2 at 37°C. To perform wound healing assay, 1.5 ×10^5^ HUVEC cells were seeded per well in 24-well plates and allowed them to adhere overnight. Before experiment, HUVECs were pretreated with or without ATN-161 (400 μM) for 12h. Sterile scratcher (SPL life sciences, 201924) was used to create a wound. Cells were washed by phosphate-buffered saline (PBS) solution, then being cultured in fetal bovine serum (FBS)-free ECM with Spp-1 (MCE,HY-P70499) (4 μg/ml) alone or combined with ATN-161 (400 μM). The inverted fluorescence microscope (Olympus IX73) was used to capture the cells at 0 and 12 h. ImageJ was used to evaluate the percentage of healing area or distance.

### Histology staining

Sirius Red Stain: 8 μm frozen sections were rehydrated with PBS for 2 min and stained according to the standard protocol of the Modified Sirius Red Staining Kit (Solarbio,20230309). Sections were imaged with an inverted fluorescence microscope (Olympus IX73).

### Quantification and statistics

All results are presented as mean ± s.e.m. Number of animals or cells represents biological replicates and is indicated in corresponding figure legends. Samples were tested using two-tailed Student’s *t* test unless otherwise indicated in figure legends. *P* values are indicated in the graphs and *P* values below 0.05 were considered to be statistically significant.

### Study approval

All animal care and experimental procedures are complied with the animal protection and welfare guidelines and were approved by Guangzhou Institutes of Biomedicine and Health, following the protocols (2023075) approved by local animal ethics committees.

### Data and Code Availability

ScRNA-seq dataset is publicly available in China National Center for Bioinformation (OMIX005618). Other data that support the findings of this study are available on reasonable request from the corresponding author.

All the analysis scripts are available from vignettes of original software webpage of Seurat and CellChat. No custom code or mathematical algorithm other than variable assignment was used in this study.

## Author Contributions

Q. Chen and Y. Liu conceived, designed, supervised the research and wrote the manuscript; R.R.G, J.W.C and Y.X.W performed most of the experiments, analyzed the data. L.Y.M performed all scRNA-seq analysis. Z.Y.Z contributed critical material. R.R.G, L.Y.M and Y.Y.L wrote the manuscript. Z.H.D, J.Z., Y.H.S and Y.Y.L performed the experiments. R.R.G, L.Y. M, J.W.C and Y.X.W share the co-first author position. Y. Liu, Q. Chen, share the co-corresponding author position.

## Acknowledgements

This work is supported by the National Key R&D Program of China (2022YFA1103200); National Natural Science Foundation of China (32270866, 32300693, 22107045); Guangzhou basic and applied basic research funding (2024A04J6259); the Fundamental Research Funds for the Central Universities (D2220650); The Pearl River Talent Recruitment Program (2021ZT09Y233); South China University of Technology (K5231040, K5220110); Talent Program and Basic Research Project of Guangzhou Institutes of Biomedicine and Health, Chinese Academy of Sciences (1103792101, GIBHBRP23-02); Chinese Academy of Medical Sciences Cancer Hospital Shenzhen Hospital Foundation (E010221005, CFA202201006); the CAS Key Laboratory of Regenerative Biology, Guangdong Provincial Key Laboratory of Stem Cell and Regenerative Medicine, Guangzhou Institutes of Biomedicine and Health, Chinese Academy of Sciences (KLRB202201); and partially supported by Science and Technology Planning Project of Guangdong Province, China (2023B1212060050). We appreciate Prof. Ralf H. Adams for kindly providing critical genetically modified mice in this study.

## References

1. Karpinska M, and Czauderna M. Pancreas-Its Functions, Disorders, and Physiological Impact on the Mammals’ Organism. Front Physiol. 2022;13:807632.

2. Storz P. Acinar cell plasticity and development of pancreatic ductal adenocarcinoma. Nat Rev Gastroenterol Hepatol. 2017;14(5):296–304.

3. Grapin-Botton A. Ductal cells of the pancreas. Int J Biochem Cell Biol. 2005;37(3):504–10.

4. Mederos MA, Reber HA, and Girgis MD. Acute Pancreatitis: A Review. JAMA. 2021;325(4):382–90.

5. Disease GBD, Injury I, and Prevalence C. Global, regional, and national incidence, prevalence, and years lived with disability for 310 diseases and injuries, 1990-2015: a systematic analysis for the Global Burden of Disease Study 2015. Lancet. 2016;388(10053):1545–602.

6. Peery AF, Crockett SD, Murphy CC, Jensen ET, Kim HP, Egberg MD, et al. Burden and Cost of Gastrointestinal, Liver, and Pancreatic Diseases in the United States: Update 2021. Gastroenterology. 2022;162(2):621–44.

7. Popa CC, Badiu DC, Rusu OC, Grigorean VT, Neagu SI, and Strugaru CR. Mortality prognostic factors in acute pancreatitis. J Med Life. 2016;9(4):413–8.

8. Kotan R, Posan J, Sapy P, Damjanovich L, and Szentkereszty Z. [Analysis of clinical course of severe acute biliary and non biliary pancreatitis: a comparative study]. Orv Hetil. 2010;151(7):265–8.

9. Ueda T, Takeyama Y, Yasuda T, Kamei K, Satoi S, Sawa H, et al. Utility of the new Japanese severity score and indications for special therapies in acute pancreatitis. J Gastroenterol. 2009;44(5):453–9.

10. Garg PK, and Singh VP. Organ Failure Due to Systemic Injury in Acute Pancreatitis. Gastroenterology. 2019;156(7):2008–23.

11. van den Berg FF, and Boermeester MA. Update on the management of acute pancreatitis. Curr Opin Crit Care. 2023;29(2):145–51.

12. Kundumadam S, Fogel EL, and Gromski MA. Gallstone pancreatitis: general clinical approach and the role of endoscopic retrograde cholangiopancreatography. Korean J Intern Med. 2021;36(1):25–31.

13. MacGoey P, Dickson EJ, and Puxty K. Management of the patient with acute pancreatitis. BJA Educ. 2019;19(8):240–5.

14. Azizoglu DB, Chong DC, Villasenor A, Magenheim J, Barry DM, Lee S, et al. Vascular development in the vertebrate pancreas. Dev Biol. 2016;420(1):67–78.

15. Cleaver O, and Dor Y. Vascular instruction of pancreas development. Development. 2012;139(16):2833–43.

16. Magenheim J, Ilovich O, Lazarus A, Klochendler A, Ziv O, Werman R, et al. Blood vessels restrain pancreas branching, differentiation and growth. Development. 2011;138(21):4743–52.

17. Chen Q, Liu Y, Jeong HW, Stehling M, Dinh VV, Zhou B, et al. Apelin(+) Endothelial Niche Cells Control Hematopoiesis and Mediate Vascular Regeneration after Myeloablative Injury. Cell Stem Cell. 2019;25(6):768–83 e6.

18. Karakatsani A, Shah B, and Ruiz de Almodovar C. Blood Vessels as Regulators of Neural Stem Cell Properties. Front Mol Neurosci. 2019;12:85.

19. Craig DJ, James AW, Wang Y, Tavian M, Crisan M, and Peault BM. Blood Vessel Resident Human Stem Cells in Health and Disease. Stem Cells Transl Med. 2022;11(1):35–43.

20. Bernier-Latmani J, Cisarovsky C, Mahfoud S, Ragusa S, Dupanloup I, Barras D, et al. Apelin-driven endothelial cell migration sustains intestinal progenitor cells and tumor growth. Nat Cardiovasc Res. 2022;1(5):476–90.

21. Liu Y, Chen Q, Jeong HW, Han D, Fabian J, Drexler HCA, et al. Dopamine signaling regulates hematopoietic stem and progenitor cell function. Blood. 2021;138(21):2051–65.

22. Liu Y, Chen Q, Jeong HW, Koh BI, Watson EC, Xu C, et al. A specialized bone marrow microenvironment for fetal haematopoiesis. Nat Commun. 2022;13(1):1327.

23. Gadomski S, Fielding C, Garcia-Garcia A, Korn C, Kapeni C, Ashraf S, et al. A cholinergic neuroskeletal interface promotes bone formation during postnatal growth and exercise. Cell Stem Cell. 2022;29(4):528–44 e9.

24. Zhao H, Huang X, Liu Z, Lai L, Sun R, Shen R, et al. Use of a dual genetic system to decipher exocrine cell fate conversions in the adult pancreas. Cell Discov. 2023;9(1):1.

25. Parte S, Nimmakayala RK, Batra SK, and Ponnusamy MP. Acinar to ductal cell trans-differentiation: A prelude to dysplasia and pancreatic ductal adenocarcinoma. Biochim Biophys Acta Rev Cancer. 2022;1877(1):188669.

26. Hyun JJ, and Lee HS. Experimental models of pancreatitis. Clin Endosc. 2014;47(3):212–6.

27. Lerch MM, and Gorelick FS. Models of acute and chronic pancreatitis. Gastroenterology. 2013;144(6):1180–93.

28. Del Poggetto E, Ho IL, Balestrieri C, Yen EY, Zhang S, Citron F, et al. Epithelial memory of inflammation limits tissue damage while promoting pancreatic tumorigenesis. Science. 2021;373(6561):eabj0486.

29. Alghamdi AAA, Benwell CJ, Atkinson SJ, Lambert J, Johnson RT, and Robinson SD. NRP2 as an Emerging Angiogenic Player; Promoting Endothelial Cell Adhesion and Migration by Regulating Recycling of alpha5 Integrin. Front Cell Dev Biol. 2020;8:395.

30. Benwell CJ, Taylor J, and Robinson SD. Endothelial neuropilin-2 influences angiogenesis by regulating actin pattern development and alpha5-integrin-p-FAK complex recruitment to assembling adhesion sites. FASEB J. 2021;35(8):e21679.

31. Wang X, Abraham S, McKenzie JAG, Jeffs N, Swire M, Tripathi VB, et al. LRG1 promotes angiogenesis by modulating endothelial TGF-beta signalling. Nature. 2013;499(7458):306-11.

32. Zhang H, Vreeken D, Junaid A, Wang G, Sol W, de Bruin RG, et al. Endothelial Semaphorin 3F Maintains Endothelial Barrier Function and Inhibits Monocyte Migration. Int J Mol Sci. 2020;21(4).

33. Jiao B, Liu S, Tan X, Lu P, Wang D, and Xu H. Class-3 semaphorins: Potent multifunctional modulators for angiogenesis-associated diseases. Biomed Pharmacother. 2021;137:111329.

34. Stoeltzing O, Liu W, Reinmuth N, Fan F, Parry GC, Parikh AA, et al. Inhibition of integrin alpha5beta1 function with a small peptide (ATN-161) plus continuous 5-FU infusion reduces colorectal liver metastases and improves survival in mice. Int J Cancer. 2003;104(4):496–503.

35. Leung CS, Yeung TL, Yip KP, Wong KK, Ho SY, Mangala LS, et al. Cancer-associated fibroblasts regulate endothelial adhesion protein LPP to promote ovarian cancer chemoresistance. J Clin Invest. 2018;128(2):589–606.

36. Berlin I, Crespo-Laumonnier B, Cournot A, Landault C, Aubin F, Legrand JC, et al. The alpha 2-adrenergic receptor antagonist yohimbine inhibits epinephrine-induced platelet aggregation in healthy subjects. Clin Pharmacol Ther. 1991;49(4):362–9.

37. Segarra M, Aburto MR, Hefendehl J, and Acker-Palmer A. Neurovascular Interactions in the Nervous System. Annu Rev Cell Dev Biol. 2019;35:615–35.

38. Wagner JUG, Tombor LS, Malacarne PF, Kettenhausen LM, Panthel J, Kujundzic H, et al. Aging impairs the neurovascular interface in the heart. Science. 2023;381(6660):897-906.

39. Sakorafas GH, Tsiotos GG, and Sarr MG. Ischemia/Reperfusion-Induced pancreatitis. Dig Surg. 2000;17(1):3–14.

40. Butler JR, Eckert GJ, Zyromski NJ, Leonardi MJ, Lillemoe KD, and Howard TJ. Natural history of pancreatitis-induced splenic vein thrombosis: a systematic review and meta-analysis of its incidence and rate of gastrointestinal bleeding. HPB (Oxford). 2011;13(12):839–45.

41. Barge JU, and Lopera JE. Vascular complications of pancreatitis: role of interventional therapy. Korean J Radiol. 2012;13 Suppl 1(Suppl 1):S45–55.

42. Wickramasinghe CU, Sivasubramanium M, and Muthugala R. Acute hemorrhagic pancreatitis following influenza infection: a case report. J Med Case Rep. 2023;17(1):176.

43. Gomez-Salinero JM, Itkin T, and Rafii S. Developmental angiocrine diversification of endothelial cells for organotypic regeneration. Dev Cell. 2021;56(22):3042–51.

44. Rafii S, Butler JM, and Ding BS. Angiocrine functions of organ-specific endothelial cells. Nature. 2016;529(7586):316–25.

45. Gomez-Salinero JM, Izzo F, Lin Y, Houghton S, Itkin T, Geng F, et al. Specification of fetal liver endothelial progenitors to functional zonated adult sinusoids requires c-Maf induction. Cell Stem Cell. 2022;29(4):593–609 e7.

46. Bar O, Gelb S, Atamny K, Anzi S, and Ben-Zvi A. Angiomodulin (IGFBP7) is a cerebral specific angiocrine factor, but is probably not a blood-brain barrier inducer. Fluids Barriers CNS. 2020;17(1):27.

47. Comazzetto S, Shen B, and Morrison SJ. Niches that regulate stem cells and hematopoiesis in adult bone marrow. Dev Cell. 2021;56(13):1848–60.

48. Puebla M, Tapia PJ, and Espinoza H. Key Role of Astrocytes in Postnatal Brain and Retinal Angiogenesis. Int J Mol Sci. 2022;23(5).

49. Gladka MM, Kohela A, Molenaar B, Versteeg D, Kooijman L, Monshouwer-Kloots J, et al. Cardiomyocytes stimulate angiogenesis after ischemic injury in a ZEB2-dependent manner. Nat Commun. 2021;12(1):84.

50. Icer MA, and Gezmen-Karadag M. The multiple functions and mechanisms of osteopontin. Clin Biochem. 2018;59:17–24.

51. Zhao H, Chen Q, Alam A, Cui J, Suen KC, Soo AP, et al. The role of osteopontin in the progression of solid organ tumour. Cell Death Dis. 2018;9(3):356.

52. Kahles F, Findeisen HM, and Bruemmer D. Osteopontin: A novel regulator at the cross roads of inflammation, obesity and diabetes. Mol Metab. 2014;3(4):384–93.

53. Sward P, Bertilsson S, Struglics A, and Kalaitzakis E. Serum Osteopontin Is Associated With Organ Failure in Patients With Acute Pancreatitis. Pancreas. 2018;47(3):e7–e10.

54. Wirestam L, Nyberg PB, Dzhendov T, Gasslander T, Sandstrom P, Sjowall C, et al. Plasma Osteopontin Reflects Tissue Damage in Acute Pancreatitis. Biomedicines. 2023;11(6).

55. Rao C, Bush N, Rana SS, Sharma RK, Rana S, Jeyashree K, et al. Plasma Osteopontin Levels as an Early Predictor of Mortality in Acute Pancreatitis: A Preliminary Study. Pancreas. 2022;51(5):e77–e9.

56. Rychlikova J, Vecka M, Jachymova M, Macasek J, Hrabak P, Zeman M, et al. Osteopontin as a discriminating marker for pancreatic cancer and chronic pancreatitis. Cancer Biomark. 2016;17(1):55–65.

57. Erikson DW, Burghardt RC, Bayless KJ, and Johnson GA. Secreted phosphoprotein 1 (SPP1, osteopontin) binds to integrin alpha v beta 6 on porcine trophectoderm cells and integrin alpha v beta 3 on uterine luminal epithelial cells, and promotes trophectoderm cell adhesion and migration. Biol Reprod. 2009;81(5):814–25.

58. Edwards DN, Salmeron K, Lukins DE, Trout AL, Fraser JF, and Bix GJ. Integrin alpha5beta1 inhibition by ATN-161 reduces neuroinflammation and is neuroprotective in ischemic stroke. J Cereb Blood Flow Metab. 2020;40(8):1695–708.

59. Zeng B, Zhou M, Wu H, and Xiong Z. SPP1 promotes ovarian cancer progression via Integrin beta1/FAK/AKT signaling pathway. Onco Targets Ther. 2018;11:1333–43.

60. Kim J, Erikson DW, Burghardt RC, Spencer TE, Wu G, Bayless KJ, et al. Secreted phosphoprotein 1 binds integrins to initiate multiple cell signaling pathways, including FRAP1/mTOR, to support attachment and force-generated migration of trophectoderm cells. Matrix Biol. 2010;29(5):369–82.

61. Beddingfield BJ, Iwanaga N, Chapagain PP, Zheng W, Roy CJ, Hu TY, et al. The Integrin Binding Peptide, ATN-161, as a Novel Therapy for SARS-CoV-2 Infection. JACC Basic Transl Sci. 2021;6(1):1–8.

62. Sui A, Zhong Y, Demetriades AM, Shen J, Su T, Yao Y, et al. ATN-161 as an Integrin alpha5beta1 Antagonist Depresses Ocular Neovascularization by Promoting New Vascular Endothelial Cell Apoptosis. Med Sci Monit. 2018;24:5860–73.

63. Cianfrocca ME, Kimmel KA, Gallo J, Cardoso T, Brown MM, Hudes G, et al. Phase 1 trial of the antiangiogenic peptide ATN-161 (Ac-PHSCN-NH(2)), a beta integrin antagonist, in patients with solid tumours. Br J Cancer. 2006;94(11):1621–6.

64. Ehling M, Adams S, Benedito R, and Adams RH. Notch controls retinal blood vessel maturation and quiescence. Development. 2013;140(14):3051–61.

65. Muzumdar MD, Tasic B, Miyamichi K, Li L, and Luo L. A global double-fluorescent Cre reporter mouse. Genesis. 2007;45(9):593–605.

66. Hao Y, Hao S, Andersen-Nissen E, Mauck WM, 3rd, Zheng S, Butler A, et al. Integrated analysis of multimodal single-cell data. Cell. 2021;184(13):3573–87 e29.

67. Bu D, Luo H, Huo P, Wang Z, Zhang S, He Z, et al. KOBAS-i: intelligent prioritization and exploratory visualization of biological functions for gene enrichment analysis. Nucleic Acids Res. 2021;49(W1):W317–W25.

68. Jin S, Guerrero-Juarez CF, Zhang L, Chang I, Ramos R, Kuan CH, et al. Inference and analysis of cell-cell communication using CellChat. Nat Commun. 2021;12(1):1088.

